# A microbiome-derived metabolite from *Staphylococcus epidermidis* inhibits *Staphylococcus aureus* biofilm formation and virulence

**DOI:** 10.64898/2026.09.23.753553

**Authors:** Ankur Sood, Kiana Hajiarbabi, Willian Rodrigues Ribeiro, Ana Carolina Oliveira, L. Caetano M. Antunes, Rosana B. R. Ferreira

## Abstract

*Staphylococcus epidermidis* is a common member of the healthy skin microbiome that contributes to barrier function and protection against pathogen colonization. In contrast, *Staphylococcus aureus* is a major human pathogen and the leading cause of skin and soft tissue infections, owing in part to its robust biofilm-forming capacity and increasing antimicrobial resistance. We previously demonstrated that *S. epidermidis* cell-free conditioned media (CFCM) inhibit *S. aureus* biofilm formation by altering bacterial gene expression without affecting growth. Here, we identify the metabolite responsible for this activity. Bioactivity-guided fractionation of *S. epidermidis* CFCM revealed pyroglutamic acid (PCA) as the active antibiofilm compound. Commercially sourced PCA inhibited *S. aureus* biofilm formation and reduced adhesion to epithelial cells. In a murine skin infection model, PCA treatment significantly attenuated disease severity, accelerated wound healing, and reduced bacterial burden. Analysis of *S. epidermidis* CFCM by chromatography, nuclear magnetic resonance spectroscopy, and mass spectrometry confirmed the presence of PCA and identified it as the bioactive constituent responsible for antibiofilm activity. Together, these findings uncover a previously unrecognized mechanism by which *S. epidermidis* suppresses *S. aureus* virulence and highlight the potential of microbiome-derived metabolites as therapeutics for the prevention and treatment of *S. aureus* skin infections.

## Introduction

The human skin serves as a critical physical barrier to the external environment, including a multitude of potential pathogens ^1^. This extensive surface is colonized by a complex and dynamic community of microorganisms known as the skin microbiota, which contributes to cutaneous homeostasis and resistance to pathogen colonization ^2^. The predominant bacterial commensals in this ecosystem include species of *Staphylococcus*, *Cutibacterium*, and *Corynebacterium* ^3^. Increasing evidence indicates that resident microbial communities actively protect the host through the production of antimicrobial and antivirulence molecules that shape microbial interactions and limit pathogen establishment ^4–13^. Despite growing recognition of the protective role of the skin microbiome, the molecular mechanisms underlying these interactions remain incompletely understood.

An example of antagonistic interaction within the skin microbiota is the competition between *Staphylococcus epidermidis*, a prevalent member of the skin microbiome, and *Staphylococcus aureus*, an opportunistic pathogen responsible for a wide range of skin and soft tissue infections ^14,15^. The success of *S. aureus* as a pathogen is closely linked to its ability to form biofilms, multicellular communities embedded within a self-produced extracellular matrix that enhances resistance to host immune defenses and antimicrobial therapies ^16,17^. *S. epidermidis* has been shown to produce several molecules that significantly inhibit *S. aureus* growth, including bacteriocins, phenol soluble modulins, and butyric acid ^6,18–20^. In contrast, relatively little is known about *S. epidermidis*-derived molecules that modulate virulence without directly affecting bacterial viability. One example, however, is the extracellular serine protease Esp, which inhibits *S. aureus* biofilm formation and nasal colonization ^21^. These observations suggest that skin commensals can regulate pathogen behavior through mechanisms that extend beyond classical antimicrobial activity.

Our previous studies identified extracellular small molecules produced by commensal *S. epidermidis* that inhibit biofilm formation and disrupt established biofilms in diverse *S. aureus* clinical isolates, including methicillin-resistant (MRSA) and susceptible (MSSA) isolates ^6,12^. These molecules also significantly altered the expression of multiple virulence-associated genes, including genes involved in biofilm development, immune evasion, quorum sensing regulation, and cytotoxins, indicating a broad impact on *S. aureus* pathogenicity. Preliminary characterization revealed that the bioactive factors present in *S. epidermidis* cell-free conditioned media (CFCM) were resistant to heat, proteases, sodium periodate, and protease inhibitor cocktails, suggesting that the activity was mediated by small, highly stable molecules distinct from Esp ^6^. However, the identity of the molecules responsible for this antivirulence activity remained unknown.

In the present study, we used bioactivity-guided fractionation coupled with spectrometric and structural analyses to identify the molecule responsible for the antibiofilm activity of *S. epidermidis* CFCM. We identified pyroglutamic acid (PCA) as a microbiome-derived metabolite that inhibits MRSA biofilm formation, reduces adherence to epithelial cells, and disrupts established biofilms. Furthermore, PCA significantly reduced bacterial burden and disease severity in a murine model of *S. aureus* skin infection. These findings uncover a previously unrecognized mechanism of interspecies interaction within the skin microbiome and demonstrate how a commensal-derived metabolite can limit *S. aureus* pathogenicity.

## Results

### Purification of bioactive molecules

To identify the molecule responsible for the antibiofilm activity present in *S. epidermidis*CFCM, we performed bioactivity-guided purification. CFCM was initially separated by reverse-phase high-performance liquid chromatography (RP-HPLC) on a C_18_ column using a methanol gradient, yielding 60 fractions. To facilitate screening, fractions were pooled and their ability to inhibit *S. aureus* biofilm formation was assessed. Among the pooled fractions, fractions 1 through 10 retained significant antibiofilm activity (**Figure 1A**). Subsequent testing of individual fractions revealed that fractions 1 and 2 accounted for the observed activity (**Figure 1B**).

**Figure 1.**
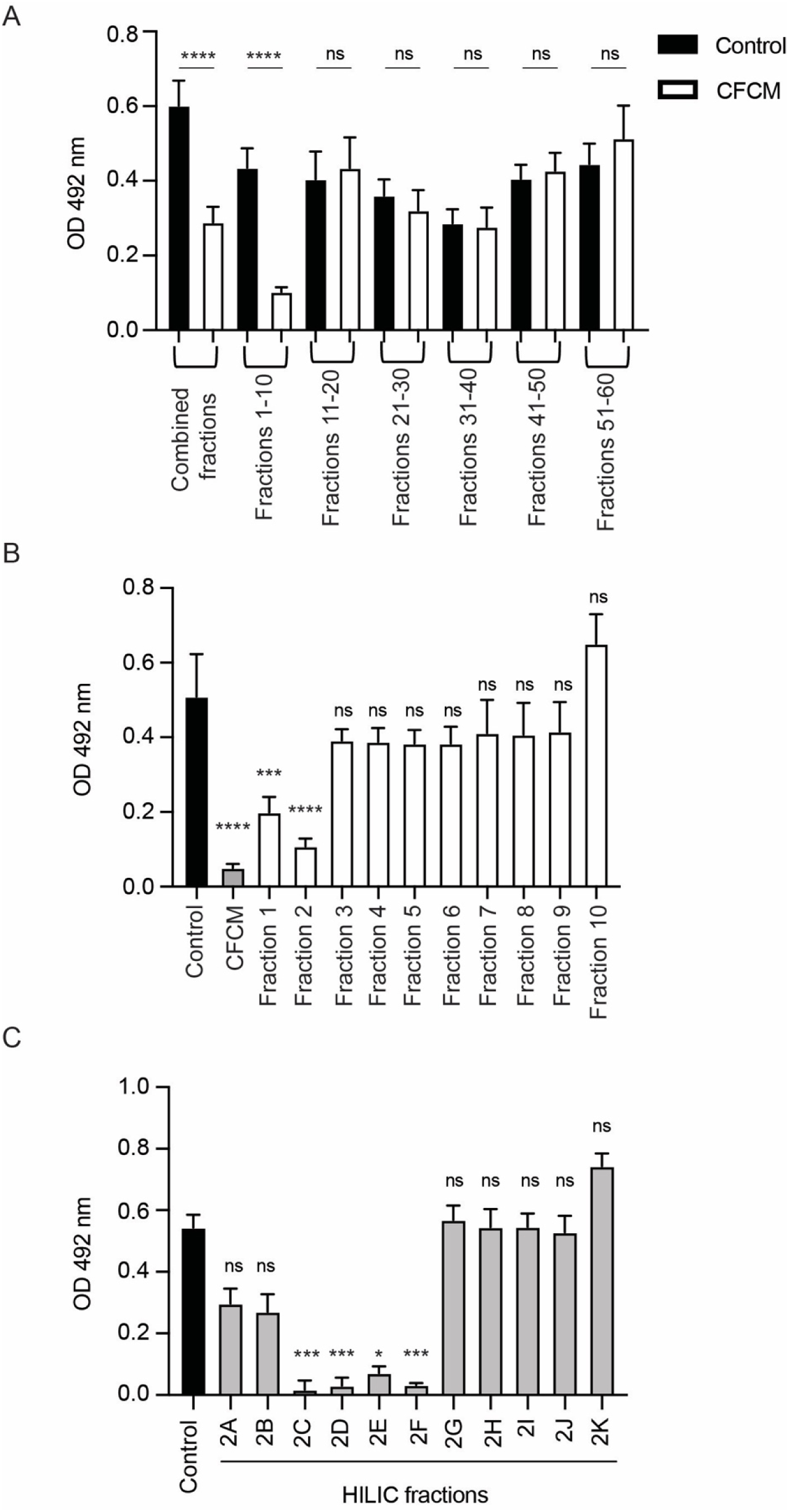
Activity-guided fractionation of *S. epidermidis* cell-free conditioned media (CFCM). A. Biofilm formation of *S. aureus* strain 1602 growing in the presence of combined fractions of control and CFCM obtained using High-Performance Liquid Chromatography (HPLC) with a C_18_ column. B. Biofilm formation of *S. aureus* strain 1602 growing in the presence of single fractions of CFCM obtained using C_18_ HPLC. C. Biofilm formation of *S. aureus* strain 1602 growing in the presence of single Hydrophilic Interaction Chromatography (HILIC) fractions obtained from HPLC fraction 2 of CFCM. Mean values of 3 independent experiments are shown, and error bars represent the SD. Statistical analyses were performed with the Kruskal-Wallis test. \**P*<0.05; \*\*\**P*<0.0001; \*\*\*\**P*<0.00001; ns: not significant.

Fraction 2 was then selected for further characterization due to its strong activity, which was comparable to that of unfractionated *S. epidermidis* CFCM. Considering that early-eluting fractions in RP-HPLC are typically enriched in polar compounds, fraction 2 was further resolved by hydrophilic interaction liquid chromatography (HILIC). Highly polar fractions cannot be effectively separated by RP-HPLC due to phase collapse and poor retention, making HILIC the appropriate alternative. This analysis generated several discrete peaks that were collected individually and evaluated for antibiofilm activity. Four fractions, designated 2C, 2D, 2E, and 2F, consistently and significantly inhibited *S. aureus* biofilm formation (**Figure 1C**), indicating that multiple metabolites contribute to the antibiofilm activity of *S. epidermidis* CFCM.

### Identification of active molecules with antibiofilm activity

To identify the bioactive molecules present in the active HILIC fractions, we performed untargeted metabolomics on fractions 2C, 2D, 2E, and 2F by ultra high-performance liquid chromatography and tandem mass spectrometry (LC-MS/MS). Inactive fractions (2H and 2K) were included as controls to facilitate the identification of metabolites enriched in the bioactive fractions. Comparative metabolomic analysis combined with database matching generated a list of 10 candidate compounds, many of which were amino acid derivatives, including valyl-proline, mebutamate, glutamyl-leucine, lysopine (N2-(1R)-1-carboxylethyl)-L-lysine), glutamyl-tyrosine, (8E)-2-amino-8-octadecene-1,2,4-triol, N6-acetyl-N6-hydroxy-lysine, pyroglutamic acid (PCA), methionylserine, and uracil (**Figure 2** and Table S1). Metabolites enriched in the highly active fraction 2F relative to an inactive fraction are shown in Table 1.

**Figure 2.**
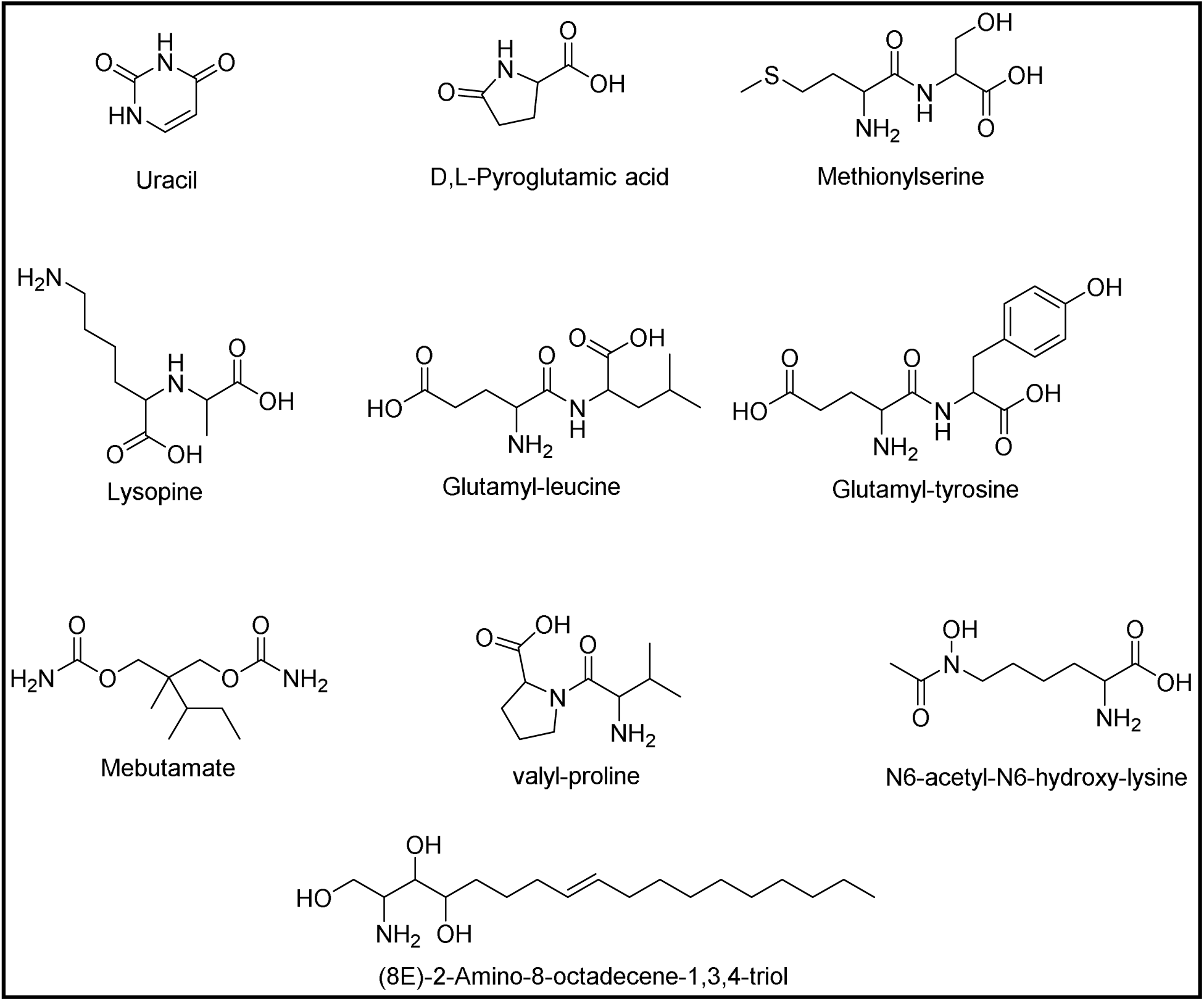
**Compounds identified in active fractions by untargeted metabolomics.**

**Table 1.** Metabolites enriched in active fraction 2F.

| Name | Formula | m/z | RT<br>[min] | Log2 Fold change |  |
| --- | --- | --- | --- | --- | --- |
|  |  |  |  | Active 2F/ Inactive<br>2H | Active 2F/ Inactive<br>2K |
| (8E)-2-amino-8-octadecene-1,3,4-triol | C<sub>18</sub>H<sub>37</sub>NO<sub>3</sub> | 338.2658 | 7.976 | 8.22 | 8.56 |
| methionylserine | C<sub>8</sub>H<sub>16</sub>N<sub>2</sub>O<sub>4</sub>S | 237.08993 | 1.548 | 5.94 | -2.11 |
| pyroglutamic acid | C<sub>5</sub>H<sub>7</sub>NO<sub>3</sub> | 130.04983 | 3.257 | 4.54 | 4.94 |
| γ-glutamyl-leucine | C<sub>11</sub>H<sub>20</sub>N<sub>2</sub>O<sub>5</sub> | 261.14375 | 3.572 | 4.13 | 5.18 |
| lysopine | C<sub>9</sub>H<sub>18</sub>N<sub>2</sub>O<sub>4</sub> | 219.13365 | 2.224 | 3.72 | 4.28 |
| γ-glutamyl-leucine | C<sub>11</sub>H<sub>20</sub>N<sub>2</sub>O<sub>5</sub> | 261.14381 | 3.441 | 3.27 | 4.48 |
| mebutamate* | C<sub>10</sub>H<sub>20</sub>N<sub>2</sub>O<sub>4</sub> | 233.1492 | 3.21 | 1.86 | 2.54 |
| valyl-proline | C<sub>10</sub>H<sub>18</sub>N<sub>2</sub>O<sub>3</sub> | 215.13861 | 3.458 | 1.83 | 2.34 |
| N6-acetyl-N6-hydroxy-lysine | C<sub>8</sub>H<sub>16</sub>N<sub>2</sub>O<sub>4</sub> | 205.11806 | 1.917 | 1.45 | 2.21 |
| γ-glutamyl-tyrosine | C<sub>14</sub>H<sub>18</sub>N<sub>2</sub>O<sub>6</sub> | 311.12309 | 3.161 | 1.42 | 1.44 |
| uracil | C<sub>4</sub>H<sub>4</sub>N<sub>2</sub>O<sub>2</sub> | 113.03467 | 2.009 | 1.09 | 1.51 |

RT: retention time; \*: not naturally found

To determine whether any of the candidate metabolites accounted for the antibiofilm activity, 7 commercially available compounds were evaluated for their ability to inhibit *S. aureus* biofilm formation. Among these, D,L-PCA emerged as the leading candidate, significantly inhibiting biofilm formation at concentrations as low as 2 µM (**Figure 3A**). Additionally, similar PCA antibiofilm activity was observed against another MRSA strain (JE2) (**Figure S1**). Uracil and valyl-L-proline also significantly reduced biofilm formation, although their profiles differ from that of PCA (**Figure 3B and C**). While PCA exhibited a monotonic dose-response, characterized by a consistent increase in the inhibitory effect proportional to the concentration used, valyl-L-proline and uracil displayed a non-monotonic dose-response, where the biological effect does not follow a linear or continuous progression with increasing dosage. In contrast, L-glutamyl-L-leucine, L-glutamyl-L-tyrosine, methionylserine, and lysopine did not significantly affect *S. aureus* biofilm formation (**Figure S2A-D**). Given its potency, reproducible dose response, and enrichment within active fractions, PCA was prioritized for further characterization as a candidate mediator of the antibiofilm activity observed in *S. epidermidis* CFCM.

**Figure 3.**
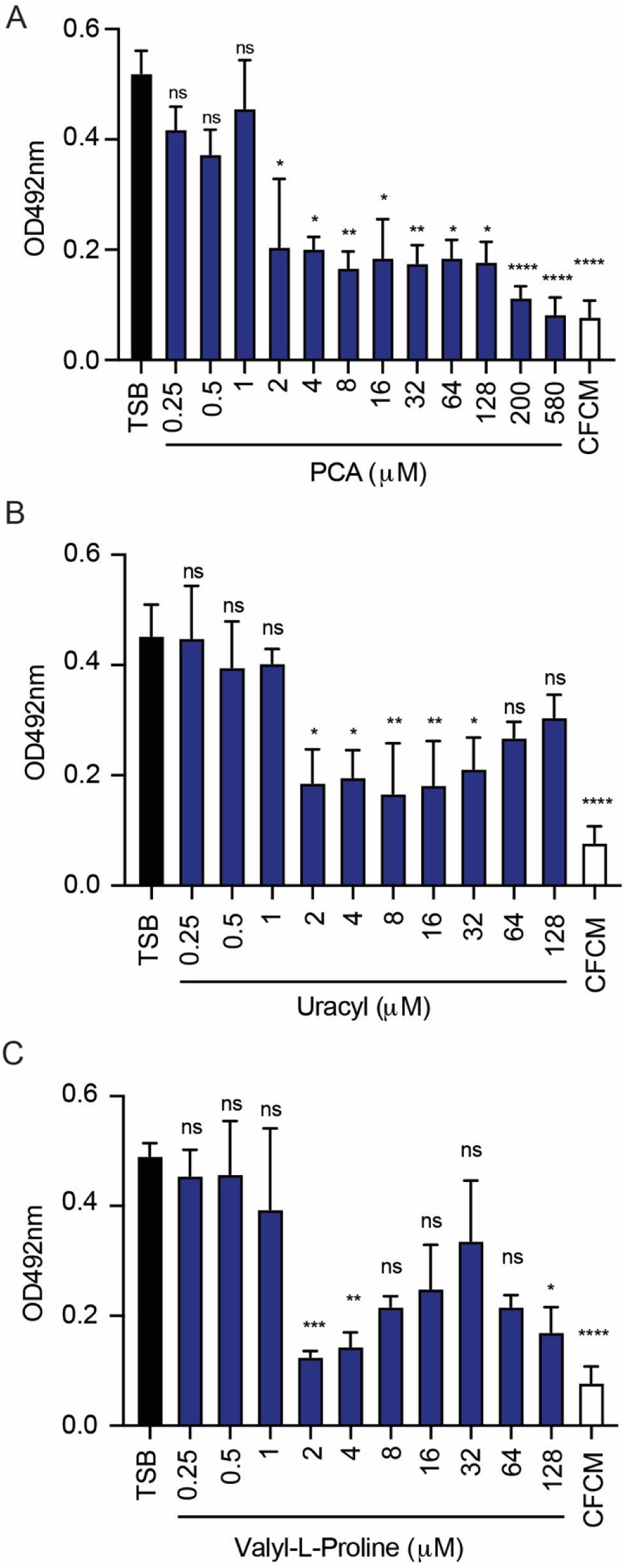
Impact of molecules identified in active fractions against *S. aureus* biofilm formation. A. Biofilm formation of *S. aureus* strain 1602 growing in the presence of different concentrations of a racemic mixture of pyroglutamic acid (PCA). B. Biofilm formation of *S. aureus* growing in the presence of different concentrations of uracil. C. Biofilm formation of *S. aureus* growing in the presence of different concentrations of valyl-L-proline. Mean values of 3 independent experiments are shown, and error bars represent the SD. Statistical analyses were performed with the Kruskal-Wallis test. \**P*<0.05; \*\**P*<0.01; \*\*\**P*<0.0001; \*\*\*\**P*<0.00001; ns: not significant.

### Effect of enantiomers of PCA

Because PCA exists as two enantiomeric forms and enantiomers can exhibit distinct biological activities ^22^, we examined whether stereochemistry influenced its antibiofilm properties. D-PCA and L-PCA were tested independently across a range of concentrations against *S. aureus* biofilm formation. Both enantiomers displayed comparable inhibitory activity, particularly at low concentrations, suggesting that, under *in-vitro* conditions, the molecular target or mechanism underlying PCA-mediated biofilm inhibition is unaffected by chirality (**Figure 4 A and B**).

**Figure 4.**
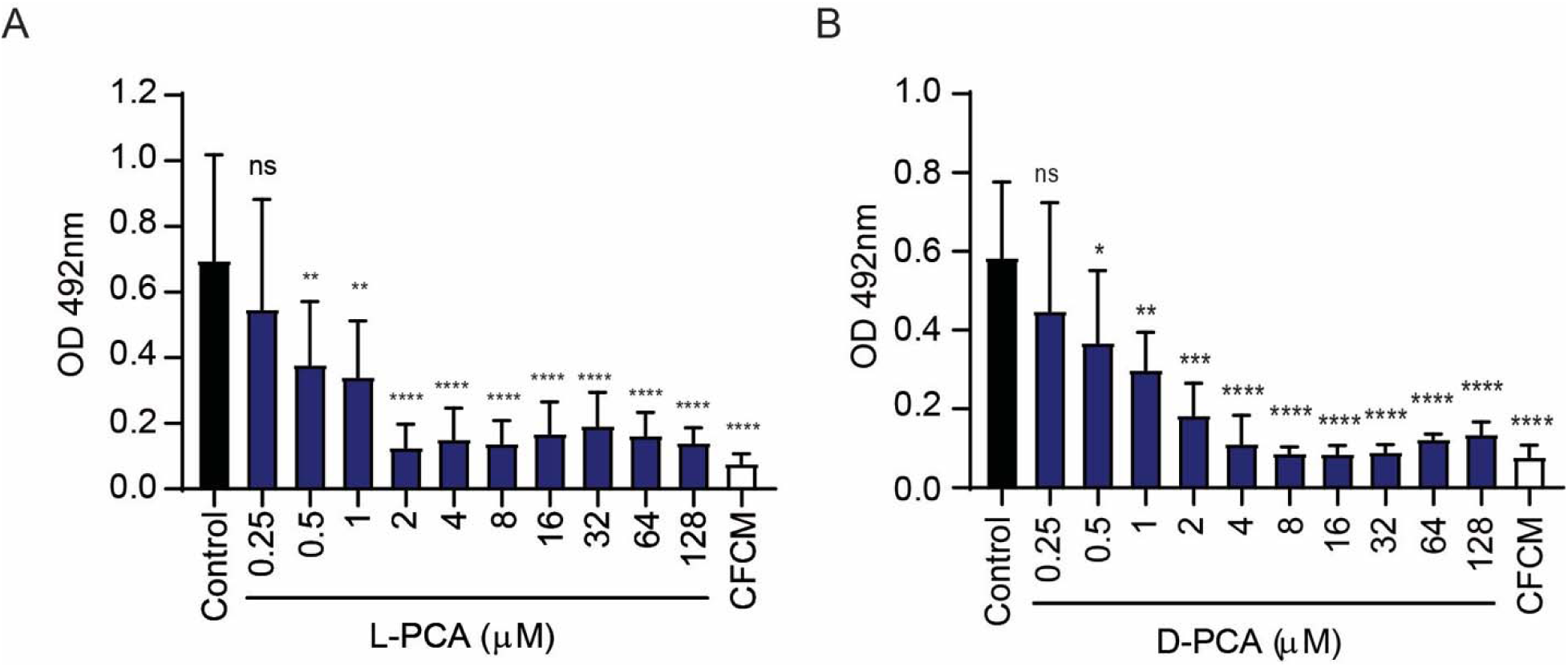
Enantiomers of pyroglutamic acid (PCA) similarly inhibit *S. aureus* biofilm formation. A. Biofilm formation of *S. aureus* strain 1602 growing in the presence of different concentrations of L-PCA. B. Biofilm formation of *S. aureus* strain 1602 growing in the presence of different concentrations of D-PCA. Mean values of 3 independent experiments are shown, and error bars represent the SD. Statistical analyses were performed with the Kruskal-Wallis test. \**P*<0.05; \*\**P*<0.01; \*\*\**P*<0.0001; \*\*\*\**P*<0.00001; ns: not significant.

### Impact of PCA on *S. aureus* planktonic growth and established biofilms

We next investigated whether PCA affected planktonic growth and established biofilms. Previously, we have shown that *S. epidermidis* CFCM does not affect *S. aureus* replication during planktonic growth ^6^. Therefore, we tested whether PCA also lacked antimicrobial activity. Consistent with what we observed with *S. epidermidis* CFCM, PCA did not impact *S. aureus* growth under planktonic conditions (**Figure 5A**). These results suggest that PCA does not impair bacterial viability and instead likely targets process involved in biofilm development and maintenance.

**Figure 5.**
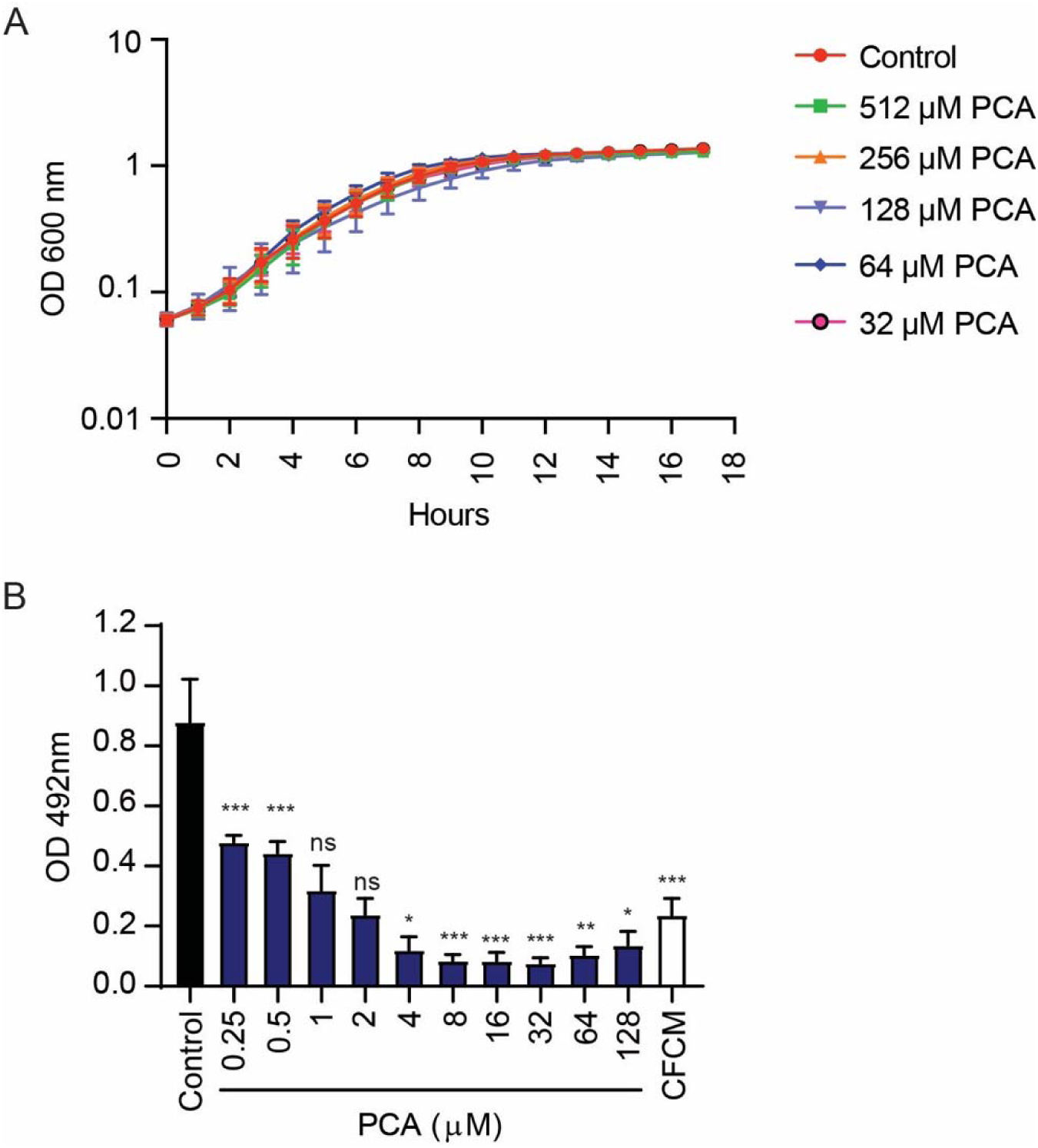
Impact of D,L pyroglutamic acid (PCA) on *S. aureus* planktonic growth and established biofilms. A. Growth curve of *S. aureus* strain 1602 in the absence or presence of different concentrations of PCA. B. Semi-quantification of *S. aureus* strain 1602 biofilm mass after treatment with either *S. epidermidis* cell-free conditioned media (CFCM) or different concentrations of PCA. Mean values of 3 independent experiments are shown, and error bars represent the SD. Statistical analyses were performed with the Kruskal-Wallis test. \**P*<0.05; \*\**P*<0.01; \*\*\**P*<0.0001; ns: not significant.

We next evaluated the impact of PCA on *S. aureus* established biofilms. Biofilms were allowed to form for 24 h before treatment with either *S. epidermidis* CFCM or PCA for an additional 24 h (**Figure 5B**). As previously observed, treatment with *S. epidermidis* CFCM significantly reduced biofilm mass relative to the control. PCA also reduced the biomass of established biofilms in a dose-dependent manner, reaching higher reduction levels than those observed with *S. epidermidis* CFCM. Together, these findings demonstrate that PCA is an active contributor to the antibiofilm properties of *S. epidermidis* and is sufficient to both inhibit biofilm formation and disrupt mature biofilms.

### Identification of PCA in *S. epidermidis* CFCM and structural confirmation

To confirm that *S. epidermidis* produces PCA, CFCM was subjected to HILIC and compared with a PCA standard. The standard eluted at a retention time of 7.2 minutes under the chromatographic conditions employed. Based on this retention profile, 7 fractions were collected from *S. epidermidis* CFCM between 3 and 8 minutes. Fractions 5, 6, and 7 (numbered by order of peaks rather than elution time), corresponding to the 6–8 min elution window, significantly inhibited biofilm formation (**Figure 6A**), consistent with the presence of one or more bioactive compounds within this region of the chromatogram.

**Figure 6.**
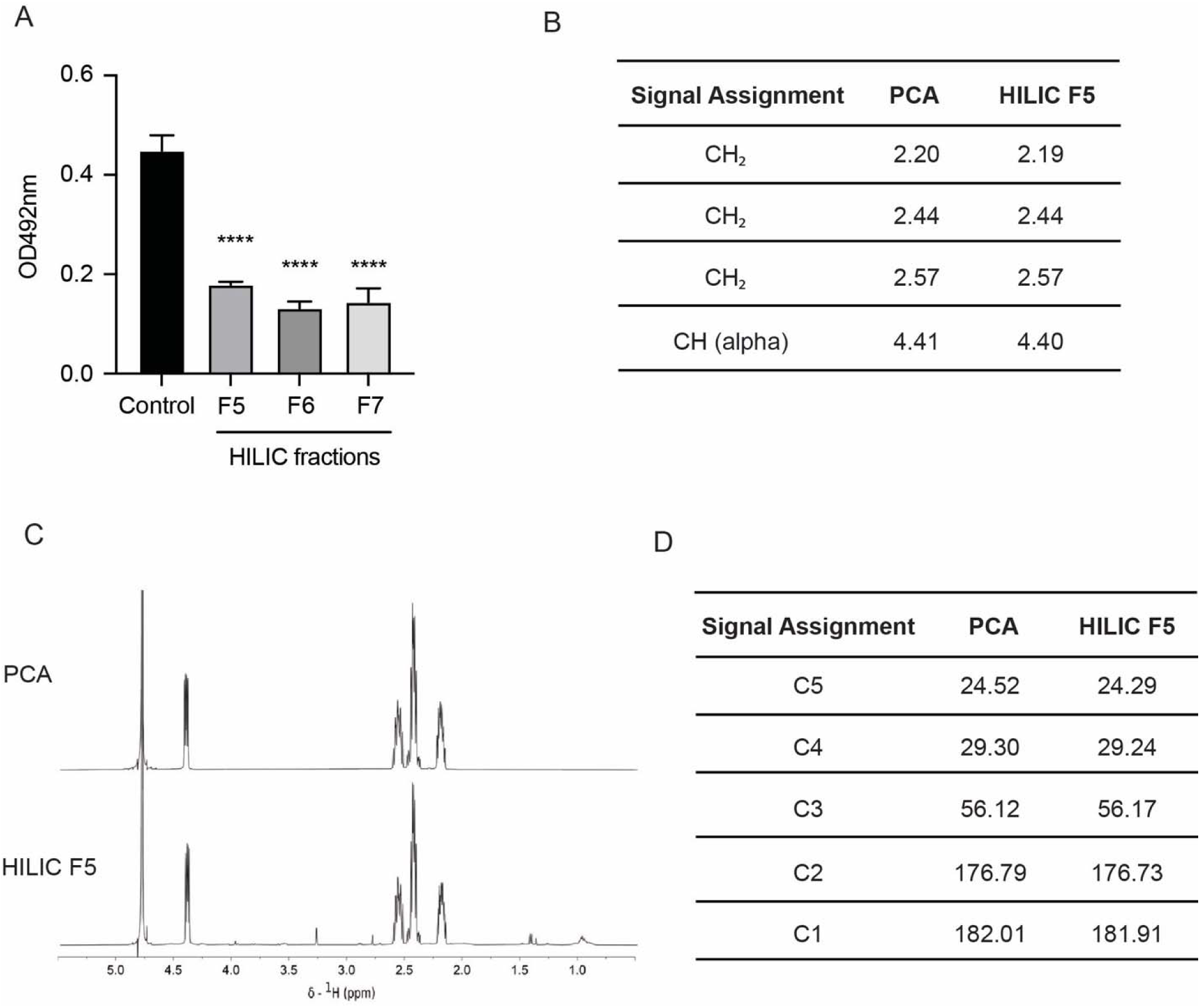
Pyroglutamic acid (PCA) is present in active fraction 5 obtained through Hydrophilic Interaction Liquid Chromatography (HILIC). A. Biofilm formation of *S. aureus* strain 1602 growing in the presence of different HILIC fractions (F5, F6, and F7). B. Comparative ^1^H Nuclear Magnetic Resonance (NMR) chemical shifts (ppm) of a PCA standard and HILIC fraction 5. C. ^1^H NMR of a PCA standard and HILIC fraction 5 dissolved in deuterium oxide (D_2_O). D. Comparative ^13^C NMR chemical shifts (ppm) of a PCA standard and HILIC fraction 5. Statistical analyses were performed with the unpaired t-test analysis. \*\*\*\**P*<0.00001.

To determine whether PCA was present in the active fractions, HILIC fraction 5 (elution time between 7.4 and 7.6), HILIC fraction 6 (elution time between 7.52 and 7.98), HILIC fraction 7 (elution time between 8.01 and 8.12) and the PCA standard were subjected to nuclear magnetic resonance (NMR) analysis. Comparison of the ^1^H NMR spectra revealed a near-identical signal pattern between active fraction 5 and the PCA standard, providing strong evidence for the presence of PCA in HILIC fraction 5 (**Figure 6B**). The same pattern was not detected in HILIC fractions 6 and 7 (data not shown). This assignment was further supported by ^13^C NMR spectroscopy, which showed agreement between the carbon chemical shifts detected in HILIC fraction 5 and those of the PCA standard (**Figure 6C and D**). The complete ^13^C NMR spectra are provided in the Supplementary Information.

To further validate the structural alignment, HILIC fraction 5 and the PCA standard were analyzed by two-dimensional NMR spectroscopy and mass spectrometry. COSY (Correlation Spectroscopy), Heteronuclear Single Quantum Coherence (HSQC), Heteronuclear Multiple Bond Correlation (HMBC) experiments revealed identical proton-carbon correlation patterns in the active fraction and the PCA standard, confirming the characteristic pyrrolidone ring structure and carbon framework of PCA (**Figures S3-6**). Mass spectrometric analysis further supported the assignment (**Figure S7**). Both samples exhibited identical parent ion masses and highly similar fragmentation patterns, providing independent confirmation of the structural assignment. Under the electrospray ionization conditions used, PCA was detected primarily as an acetonitrile adduct ion, a common phenomenon for small polar metabolites analyzed in acetonitrile-containing solvent systems. The observed mass was consistent with the expected molecular composition of PCA and its corresponding adduct species. Collectively, the concordance of chromatographic behavior, 1D and 2D NMR data, and mass spectrometric analyses conclusively identified PCA as a constituent of *S. epidermidis* CFCM and support its role as a bioactive metabolite contributing to the antibiofilm activity of the conditioned medium.

### Impact of PCA on *S. aureus* adhesion to host cells and subcutaneous infection

Because bacterial adhesion to host tissues is a critical early step in *S. aureus* colonization and biofilm development in biotic surfaces, we next evaluated the effect of PCA on bacterial attachment to epithelial cells. Before doing so, we sought to determine whether PCA exerted toxic effects on host cells. A549 cell viability was assessed following treatment with the concentrations used throughout the study, and the results showed that PCA did not significantly affect epithelial cell viability (**Figure S8**). However, exposure to PCA significantly reduced *S. aureus* adhesion to A549 epithelial cells compared with untreated controls (**Figure 7A**), indicating that the effects of PCA extend beyond inhibition of biofilm formation on abiotic surfaces and influence biologically relevant host-pathogen interactions.

**Figure 7.**
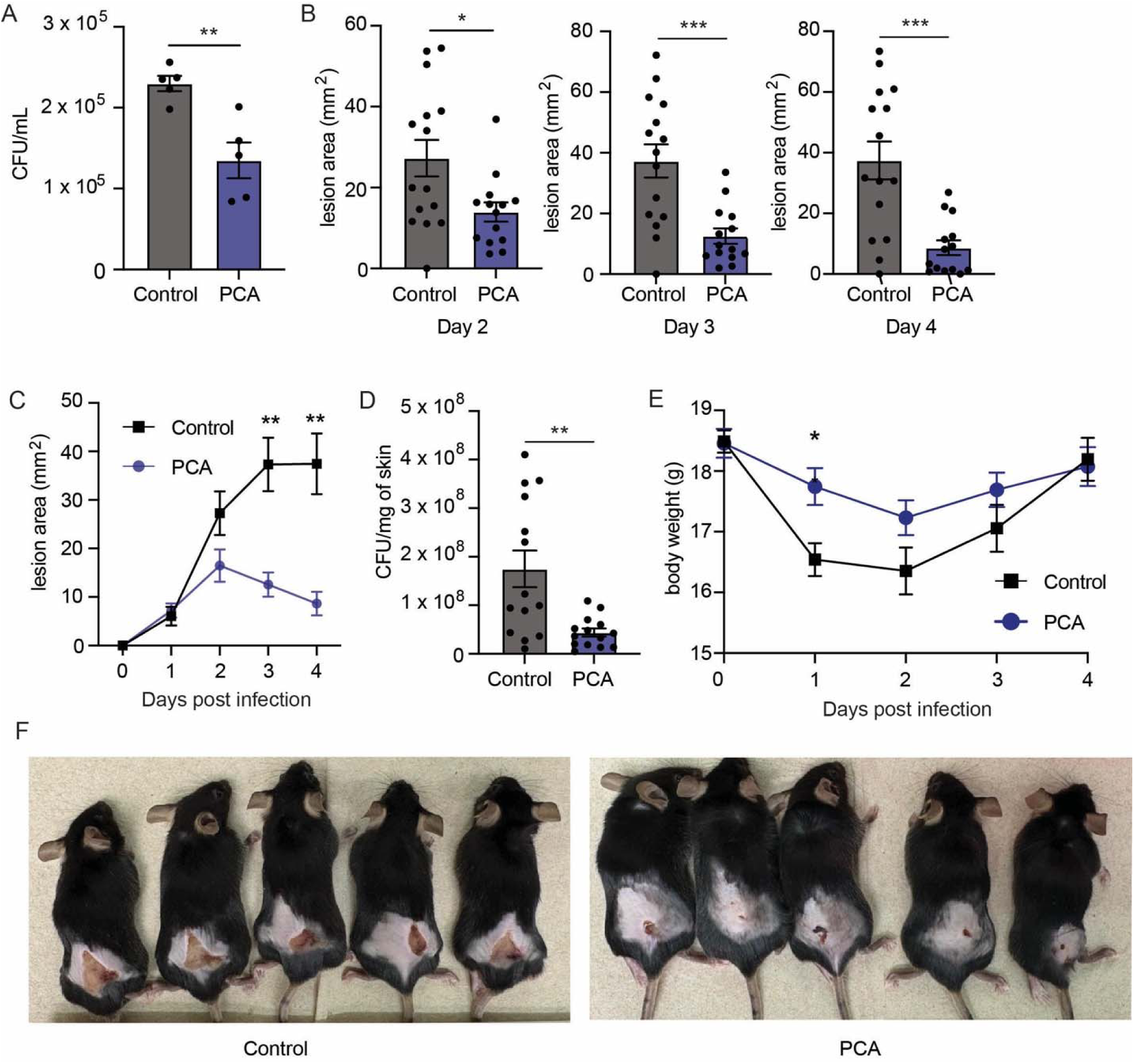
Pyroglutamic acid (PCA) reduces *S. aureus* adhesion to A549 epithelial cells and attenuates disease severity in a murine subcutaneous skin infection model. A. Impact of D,L-PCA (200 µM) on *S. aureus* strain JE2 adhesion to A549 cells. B. Skin lesion sizes 2, 3, and 4 days post infection of mice treated with vehicle (water) or PCA. C. Wound kinetics in mice treated with vehicle or PCA. D. *S. aureus* colony-forming units (CFU) recovered from the skin lesion of mice treated with vehicle or PCA. E. Body weight of mice treated with vehicle or PCA over 4 days of infection. F. Representative images at 4 days post-infection of mice treated with vehicle or PCA. For A, dots represent CFU/mL values obtained for each well and bars represent SEM. For B-D, dots represent values obtained for each mouse, and bars represent SEM. * *P*<0.05, ** *P<*0.01, *** *P*<0.001 using Welch’s t-test (A, B and D), Two-way mixed ANOVA (C), and Two-way ANOVA (E).

We next investigated the impact of PCA in a murine model of subcutaneous *S. aureus* infection. Mice were infected with *S. aureus* cultured in the presence or absence of PCA and subsequently treated daily with PCA or vehicle. PCA treatment significantly reduced lesion size, accelerated wound resolution, and decreased bacterial burdens within infected skin tissues (**Figures 7B-D**). PCA-treated mice also experienced less infection-associated weight loss than control animals (**Figure 7E)**. Representative lesion images confirmed the reduced severity of infection in PCA-treated animals (**Figure 7F**). Notably, no adverse effects were observed in uninfected mice receiving PCA. Collectively, these findings demonstrate that PCA is effective in reducing both disease severity and bacterial persistence during *S. aureus* skin infection *in vivo*.

## Discussion

The skin microbiome plays a critical role in protecting the host from pathogen colonization through a combination of immune modulation, niche competition, and the production of bioactive molecules that influence pathogen behavior ^3^. Although numerous studies have documented antagonistic interactions between commensal and pathogenic microorganisms, the specific metabolites that mediate these effects remain incompletely characterized ^8,14,23^. In this study, we identified PCA as a bioactive metabolite produced by *S. epidermidis* that suppresses multiple virulence-associated phenotypes of *S. aureus*. Through activity-guided fractionation, structural characterization, and functional analyses, we demonstrate that PCA inhibits biofilm formation, disrupts established biofilms, reduces epithelial cell adhesion, and limits pathogenicity *in vivo*, all without affecting planktonic bacterial growth.

*S. epidermidis* is known to produce various metabolites that modulate the growth and virulence of *S. aureus* ^18–21,24^. Previous work from our group demonstrated that extracellular factors present in *S. epidermidis* CFCM inhibit biofilm formation in diverse *S. aureus* clinical isolates, including both MRSA and MSSA strains ^6^. These molecules were also able to disrupt established biofilms of *S. aureus* strains, and the effect was observed in strains producing both protein and polysaccharide-based biofilms. Additionally, *S. epidermidis* CFCM significantly decreased the concentration of antibiotic necessary to eradicate biofilms. However, the identity of the bioactive molecules responsible for these effects remained unknown. The present study advances those findings by identifying PCA as one of the main constituents of the active fraction and establishing its contribution to the antivirulence activity of *S. epidermidis*. Notably, PCA recapitulated several phenotypes previously observed with conditioned medium, including inhibition of biofilm formation and disruption of mature biofilms, supporting its role as an important mediator of interspecies interactions within the skin microbiome.

A notable feature of PCA is its lack of detectable effects on planktonic growth. Unlike conventional antimicrobials, which impair bacterial viability and therefore impose strong selective pressure for resistance, antivirulence molecules target pathogenic behaviors that are required for successful host colonization and disease progression ^25^. By inhibiting biofilm formation, reducing host-cell adhesion, and limiting skin infection without affecting bacterial replication, PCA displays characteristics consistent with this emerging class of therapeutic agents. Although antivirulence molecules have not yet been approved for clinical use, they represent a potential breakthrough strategy to combat antimicrobial-resistant pathogens. Combining antivirulence compounds with existing antibiotics may enhance their antimicrobial activity, reducing the doses needed and extending their use.

PCA, also known as 5-oxoproline, is a naturally occurring cyclic derivative of the amino acid glutamic acid that is widely distributed in biological systems ^26^. Previous studies have reported that glutamic acid, the precursor of PCA, has antibiofilm activity against *S. aureus* ^27^. However, this effect required substantially higher concentrations than those observed in the present study. Whereas D-glutamic acid has been reported to exert modest antibiofilm activity at millimolar concentrations (40 mM), PCA significantly inhibited biofilm formation at low micromolar concentrations. This marked difference in potency suggests that cyclization of glutamic acid may substantially enhance biological activity. Furthermore, the lack of stereospecificity between D- and L-PCA in the present study suggests that PCA does not act through a highly stereoselective interaction with a protein target, potentially indicating a more general physiological or regulatory mechanism. Whether PCA is generated enzymatically by *S. epidermidis* or arises through spontaneous cyclization remains unknown, but our findings indicate that the cyclic derivative possesses distinct functional properties that warrant further investigation.

In addition to its newly identified role as an antivirulence metabolite, PCA has established biological relevance within the skin environment. PCA is a major component of the natural moisturizing factor (NMF) and contributes to skin hydration and barrier function ^28^. It is intriguing that PCA may contribute both to host barrier function and to microbiome-mediated colonization resistance, suggesting that this metabolite occupies a unique position at the interface between host physiology and microbial ecology. Interestingly, reduced levels of PCA have been reported in psoriatic lesions, conditions that are frequently associated with increased *S. aureus* colonization ^29–31^. Although the relationship between these observations remains speculative, they raise the possibility that PCA may contribute to colonization resistance within the skin ecosystem. Future studies investigating the association between cutaneous PCA levels, microbiome composition, and susceptibility to *S. aureus* colonization may provide additional insight into the ecological significance of this metabolite.

Untargeted metabolomic analysis further identified uracil and valyl-proline as candidate metabolites with antibiofilm activity. In contrast to PCA, however, both compounds exhibited non-monotonic dose-response relationships. Such biphasic responses have been described in diverse biological systems and can arise through multiple mechanisms, including alterations in target engagement, signaling pathways, or metabolic adaptation ^32^. While these findings suggest that additional metabolites may contribute to the antibiofilm activity of *S. epidermidis* conditioned medium, their biological roles and mechanisms of action remain to be determined. Nevertheless, the identification of additional active metabolites suggests that the antibiofilm activity of *S. epidermidis* CFCM results from a chemically diverse repertoire of molecules that may act additively or synergistically.

The structural assignment of PCA was supported by multiple independent analytical approaches. Concordant chromatographic retention times, highly similar mass spectrometric profiles, and matching ^1^H and ^13^C NMR spectra between the active fraction and the PCA standard provided strong evidence for its identity. The minor deviations are well within the standard range of experimental variance, often caused by slight differences in sample concentration ^33^. This assignment was further confirmed through COSY, HSQC, and HMBC analyses, which established the expected proton-proton and proton-carbon correlations characteristic of PCA. Together, these data conclusively identify PCA as a component of the bioactive fractions derived from *S. epidermidis* conditioned medium. The observation that adjacent HILIC fractions also retained antibiofilm activity suggests either chromatographic tailing of PCA or the presence of additional bioactive metabolites that remain to be identified.

Despite the identification of PCA as a bioactive constituent of *S. epidermidis* conditioned medium, several important questions remain unanswered. First, the biosynthetic origin of PCA in *S. epidermidis* is currently unclear. Unlike thermophilic lactic acid bacteria reported to produce PCA through glutamine-to-PCA cyclization activity ^34^, *S. epidermidis* has not, to our knowledge, been reported to encode or use a characterized dedicated PCA-producing enzyme. Its annotated 5-oxoproline-associated genes, including *pxpABC* and *pcp*, are more consistent with PCA turnover and peptide processing than with a defined biosynthetic pathway ^35,36^. A dedicated enzymatic PCA-biosynthetic pathway has not been characterized in *S. epidermidis*. PCA may therefore arise through previously uncharacterized enzymatic pathways or through spontaneous cyclization of glutamic acid and related intermediates ^34,37^. Defining the molecular basis of PCA production will be important for understanding how its synthesis is regulated within the skin environment. A second major question that remains is regarding the mechanism by which PCA attenuates *S. aureus* virulence. Our previous studies demonstrated that *S. epidermidis* conditioned medium alters the expression of multiple *S. aureus* virulence-associated genes, including key transcriptional regulators ^6^. It is therefore possible that PCA contributes to these transcriptional changes, either directly or indirectly. Alternatively, PCA may interfere with metabolic pathways, surface-associated structures, or regulatory networks required for biofilm formation and host interaction. Future transcriptomic or proteomic analyses will be required to determine whether PCA directly modulates regulatory pathways governing biofilm development and virulence. Elucidating the molecular targets of PCA will be necessary to fully understand how this metabolite influences *S. aureus* physiology and pathogenicity.

The *in vivo* findings further support the biological relevance of PCA-mediated inhibition of *S. aureus* virulence. PCA treatment significantly reduced lesion size, accelerated wound resolution, and decreased bacterial burdens in a murine skin infection model. These beneficial effects were accompanied by reduced infection-associated weight loss and occurred without detectable toxicity. The concordance between the *in vitro* and *in vivo* data suggests that interference with early colonization and biofilm-associated processes translates into measurable reductions in disease severity. Importantly, the reduction in bacterial burden indicates that modulation of virulence traits *in vitro*, such as biofilm formation and host cell adhesion, can have meaningful consequences for pathogen persistence during infection. Additional studies should focus on determining additional factors affected in *S. aureus* responsible for the PCA-mediated phenotypes *in vivo*. In addition, the contribution of host immune responses to the protective effects observed *in vivo* was not examined.

In conclusion, this study identifies PCA as a previously unrecognized microbiome-derived metabolite that attenuates multiple virulence-associated traits of *S. aureus*. PCA inhibited biofilm formation, disrupted established biofilms, reduced epithelial cell adhesion, and attenuated mouse skin infection, without affecting bacterial growth *in vitro*. These findings reveal a novel mechanism through which *S. epidermidis* potentially contributes to colonization resistance and highlight the value of microbiome-derived metabolites as sources of antivirulence therapeutics. More broadly, our results underscore the importance of understanding metabolite-mediated interactions within the skin microbiome as a foundation for the development of new strategies to combat antimicrobial-resistant pathogens.

## Methods

### Bacterial strains and growth conditions

The bacterial strains used in this study have been described previously ^6^. The commensal *S. epidermidis* RF1 isolate was recovered from healthy human skin and served as the source of CFCM. MRSA isolates (1602 and JE2) were used throughout the study. Strain 1602 is a clinical isolate ^6^, whereas JE2 is lab derivative of the USA300 lineage ^38^. Bacterial stocks were maintained at -80 °C in trypticase soy broth (TSB; BD Diagnostics, Maryland, USA) supplemented with 20% glycerol (Fisher Bioreagents, Waltham, USA). For routine culture, bacteria were streaked onto trypticase soy agar (TSA; BD Diagnostics) plates and incubated at 37 °C for 24 h.

### Preparation of *S. epidermidis* cell-free conditioned media (CFCM)

*S. epidermidis* RF1 was cultured in 50 mL of TSB for 24 h at 37 °C with shaking (250 rpm). Cultures were centrifuged at 3,100 ×*g* for 15 min and supernatants sterilized through 0.22-μm membrane filters. CFCM were concentrated using a Speed Vac Concentrator plus (Savant SPD140DDA SpeedVac concentrator; Thermo Fisher Scientific, Waltham, USA) and the dried material was resuspended in sterile distilled water to yield a 20-fold concentrate. CFCM was added to assays at a final concentration equivalent to 2X the original supernatant. Media processed identically in the absence of bacterial growth were used as controls ^6^.

### Biofilm formation assays

Biofilm formation was quantified using a microtiter plate assay, as previously described^6^. Briefly, bacterial colonies were used to prepare cell suspensions in sterile distilled water (OD_600nm_ 0.1) and inoculated (10% v/v) into TSB supplemented with 1% glucose containing either CFCM, PCA, other test compounds, or controls (vehicles). Cultures were incubated for 24 h at 37 °C in 96-well polystyrene plates (TPP, Trasadingen, Switzerland). After incubation, wells were washed three times with phosphate buffered saline (PBS pH 7.2; Thermo Fisher Scientific), dried at 60 °C for 1 h and stained with a 0.1% safranin solution (Sigma-Aldrich, St. Louis, USA) for 15 min. Excess stain was removed by rinsing the wells twice with PBS and the stain bound to biofilms was solubilized in a 95% ethanol solution. After 30 min, biofilm biomass was quantified by measuring absorbance at 492nm (OD_492nm_) using a multimode microplate reader (VaroskanLux, Thermo Fisher Scientific). Values were normalized against the average ODs of the negative controls (media-only). Experiments were performed using three biological replicates and repeated at least three independent times.

### Screening of candidate metabolites and PCA enantiomers

Commercial standards corresponding to candidate metabolites identified by metabolomic analysis were obtained and tested for inhibition of *S. aureus* biofilm formation using the assay described above. D, L-pyroglutamic acid (PCA) (TCI, Maryland, USA, CAS: 149-87-1), uracil (TCI, Maryland, USA, CAS: 66-22-8), valyl-L-proline (Synthesized by Synthetic Chemical Biology Core Facility at KU), L-glutamyl-L-leucine (AA Blocks Inc., San Diego, USA, CAS:2566-39-4), L-glutamyl-L-tyrosine (A2B Chem, San Diego, USA, CAS:7432-23-7) methionylserine (Synthesized by Synthetic Chemical Biology Core Facility at KU), and lysopine (Synthesized by Synthetic Chemical Biology Core Facility at KU) were evaluated across a range of concentrations. To evaluate the effect of stereochemistry, D-PCA (TCI, Maryland, USA, CAS: 4042-36-8) and L-PCA (Sigma Aldrich, St Louis, USA, CAS: 98-79-3) were tested independently using identical biofilm formation assays over the concentration ranges indicated in the figures.

### Planktonic growth assays

The effect of PCA on bacterial growth was assessed in 96-well plates (TPP). Overnight cultures of *S. aureus* in TSB were diluted to an initial OD_600nm_ of 0.05 in TSB containing the indicated concentrations of PCA. Plates were incubated at 37 °C with continuous shaking (120 rpm) and the OD_600nm_ was measured hourly using a microplate reader (SpectraMax, Molecular Devices, San Jose, CA, USA).

### Treatment of established biofilms

*S. aureus* biofilms were established in TSB supplemented with 1% glucose for 24 h before treatment, as described earlier ^6^. Following two washes with sterile PBS, wells received either CFCM, PCA at the indicated concentrations, or vehicle control (sterile PBS). Plates were incubated for an additional 24 h at 37 °C. Biofilm biomass was then quantified using the safranin staining procedure described earlier. Experiments were performed in triplicate at two independent times.

### Host cell adhesion assays

A549 lung epithelial cells (5×10^4^ cells/well; ATCC, Manassas, USA) were maintained in high-glucose Dulbecco’s Modified Eagle Medium (DMEM; Thermo Fisher Scientific) supplemented with 10% heat-inactivated fetal bovine serum (FBS; Cytiva, Marlborough, USA) at 37 °C in 5% CO_2_. For the assays, cells were plated in 24-well polystyrene plates (1×10^5^ cells/well) and incubated at 37 °C in a CO_2_ atmosphere for 24 h. After incubation, the media was discarded, the cells were washed with PBS, and fresh DMEM medium containing 10% heat inactivated FBS was added 1 h prior to infection. *S. aureus* overnight cultures in TSB were subcultured to OD_600nm_ 0.05 in the presence or absence of 200 μM PCA until they reached OD_600nm_ 0.4 (mid-logarithmic phase) at 37 °C (250 rpm). Then, cultures were centrifuged (13,000×g for 5 min), and the pellet washed with PBS (Corning Inc., Corning, USA). The cell pellet was then resuspended in DMEM, and A549 cells were infected at a MOI of 25. The plates were incubated at 37 °C in 5% CO_2_ for 1 h. Loosely bound bacteria were removed from the cell monolayers by two washes with PBS. The cells were then lysed with 0.1% Triton X 100 (Thermo Fisher Scientific) and serial dilutions were plated on TSA agar to determine total viable bacteria. Each experiment was performed with 5 replicates in at least two independent days.

### Cell viability assays

The cytotoxicity of PCA toward A549 cells was evaluated using an MTT (3-(4,5-dimethylthiazol-2-yl)-2,5-diphenyltetrazolium bromide) assay. Cells were seeded in 96-well plates (1×10^5^ cells/well) in DMEM and incubated overnight at 37 °C and 5% CO_2_ to allow cell attachment. After incubation, the cells were exposed to PCA (10, 50, 100, and 200 µM) or vehicle, prepared in DMEM (Thermo Fisher Scientific) supplemented with 10% heat-inactivated FBS (Cytiva), and incubated for 22 h. MTT reagent (5 mg/mL; Sigma Aldrich, Milwaukee, USA) was then added to a final concentration of 0.5 mg/mL and cells were incubated for an additional 2 h at 37 °C and 5% CO_2_. The plates were then centrifuged at 200×g for 5 minutes and the supernatants carefully discarded. Formazan crystals were dissolved in dimethyl sulfoxide (DMSO; Sigma Aldrich, Milwaukee, USA) and absorbance was measured at 570 nm. Cells treated with PBS (1X) served as the negative viability control, while cells maintained in DMEM with 10% FBS served as the positive control. All treatments were performed in quintuplicates. Cell viability was expressed as a percentage relative to the positive control group ^39^.

### RP-HPLC fractionation

Bioactive compounds present in *S. epidermidis* CFCM were isolated using preparative reverse-phase high-performance liquid chromatography (RP-HPLC). Concentrated CFCM was dissolved in 20% methanol, filtered through a 0.22-µm membrane (PES), and separated on a Prep C_18_ column (5 µM OBD, 19×250 mm, Waters, Milford, USA). Elution was performed using a water-methanol gradient containing 0.1% trifluoroacetic acid (TFA). The gradient ran as follows: 0-10 mins – 95-90% water; 11-55 mins – 90-50% water; 55-65 min – 50-10% water. Flow rate was 12 mL/min and fractions were collected at one-minute intervals for 60 min, dried under nitrogen, resuspended in distilled water, and screened for antibiofilm activity against *S. aureus* 1602. The same steps were followed for the control supernatant.

### HILIC fractionation

Active RP-HPLC fraction 2 or unfractionated CFCM were further separated by hydrophilic interaction liquid chromatography (HILIC). Samples were dried and dissolved in 95% acetonitrile supplemented with 2-3 drops of ethanol to complete the solubilization and injected onto a BEH Z-HILIC column (5µM OBD^TM^ 19 × 250 mm, Waters). The mobile phase consisted of 95% acetonitrile and 5% water containing 0.1% TFA. Fractions corresponding to individual peaks were collected, dried under nitrogen, dissolved in distilled water, and evaluated for antibiofilm activity against *S. aureus* 1602.

### Untargeted metabolomic analysis

Active (2C-2F) and inactive (2H and 2K) HILIC fractions were analyzed using a Vanquish HPLC system coupled with a Q-Exactive HF (QE-HF) mass spectrometer (ThermoFisher Scientific). Chromatographic separation was performed using a Hypersil GOLD™ column; 2.1 mm x 150 mm, 1.9 µm; (ThermoFisher Scientific) operating at a column temperature of 40C. Elution was performed using a15 minutes reverse phase (RP) gradient at a flow rate of 0.3mL/min using two solvents: Solvent A consisting of 0.1% formic acid in water and solvent B consisting of 0.1% formic acid in methanol. Sample vials were stored at 4C in the HPLC autosampler, and 3µL were injected both for positive and negative ionization mode. Samples were analyzed separately both in MS and MS/MS mode. For MS analysis, QE-HF resolution of 120,000 was used and for MS/MS analysis a resolution of 60,000 was used. Blank samples were injected at the beginning and end of each batch; QC samples were also analyzed after every 8 injections to assess system stability throughout the batch. Raw instrument data (RAW) were exported and analyzed using Compound Discoverer 3.1, mass accuracy was set to 5ppm. Metabolic features enriched in active fractions were identified by comparative analyses.

Candidate metabolites were prioritized based on enrichment in active fractions and commercial availability for functional testing.

### NMR and mass spectrometric analyses

Active HILIC fractions and PCA standards were dissolved in deuterium oxide (99.8 atom % D, Thermo Scientific) and analyzed at the University of Kansas NMR Core Facility using a Bruker AVIII 500 MHz spectrometer equipped with a CPUL cryoprobe and CASE autosampler. One-dimensional ^1^H and ^13^C NMR spectra as well as COSY, HSQC, and HMBC experiments were acquired using standard pulse sequences.

For mass spectrometry, HILIC fraction 5 and PCA standard were dissolved in distilled water containing a few drops of methanol and analyzed at the University of Kansas Synthetic Chemical Biology Core Facility using electrospray ionization liquid chromatography-mass spectrometry (LC-ESI-MS). Structural assignments were based on comparison of chromatographic behavior, mass spectra, and NMR data with PCA standards.

### Murine subcutaneous infection model

Animal studies were conducted in strict accordance with a protocol (293-01) approved by the University of Kansas Institutional Animal Care and Use Committee and NIH guidelines. Four days prior to infection, 7-week-old C57BL/6J female mice (Jackson Laboratories, Bar Harbor, USA) were anesthetized with ketamine and xylazine (87.5 mg/kg and 12.5 mg/kg, respectively), and the hair on their lower back removed using a veterinary shaver, followed by a depilatory cream (Nair, Church & Dwight, Ewing, USA). The cream was removed with isopropanol wipes, and the area was treated with ointment (Aquaphor, Beiersdorf, Inc., Wilton, USA). Aquaphor was reapplied to the area daily for the following two days. On the day of infection, an overnight culture of *S. aureus* JE2 was diluted 1:100 into 10 mL TSB in a 50-mL Falcon tube and grown in the absence or presence of PCA (200 µM) for approximately 3 h (OD_600nm_ of 1.4). Cultures were then centrifuged at 1,730 ×*g* for 10 minutes, and the supernatants were removed. The bacterial pellets were washed with 1X PBS and centrifuged again. The resulting pellets were resuspended in either 1X PBS (control) or PBS containing 200 µM PCA (treated) and adjusted to an OD_600nm_ of 1.4. Mice were then inoculated subcutaneously with approximately 6×10^7^ CFU of *S. aureus* grown with or without PCA (100 µL) into their lower back (right-side) using a 27-gauge needle. For the following three days, mice were injected subcutaneously at the site of infection with either 100 µL of PCA (1.73 mM) or water (control) daily. Mice body weight and skin lesion size were measured and recorded daily. After 4 days of infection, mice were euthanized using CO_2_ asphyxiation and cervical dislocation as the secondary method. Skin tissues were collected into 2-mL lysing matrix H tubes (MP Biomedicals, Irvine, USA) containing buffer (1X HBSS with 0.2% HAS and 10 mM HEPES). Tissues were homogenized using a Tissuelyzer III (Qiagen, Germantown, USA) using 5 cycles of 1 min each at 30 Hz with 1 min of cooling in ice between cycles. Bacterial loads on skin (site of injection) were determined through serial dilution in PBS containing 0.1% Triton for the first dilution to minimize bacterial aggregation. Dilutions were plated in duplicate in TSA and incubated for 24 h at 37 °C for colony-forming units (CFU) enumeration. The skin lesion area was determined using ImageJ software (NIH). Uninfected control mice treated with PCA or vehicle (n=5 each) were included to ensure that no lesions or CFUs were detected under the conditions used. Two independent experiments were performed, comprising a total of 15 animals per group.

### Statistical analyses

Statistical analyses were performed using GraphPad Prism, version 11.0.0. The statistical tests used for each experiment are indicated in the corresponding figure legends. Differences were considered statistically significant when values of *P <* 0.05.

## Data availability

The datasets generated during this study are available from the corresponding author upon reasonable request.

## Acknowledgements

This study was supported by startup funds provided by the University of Kansas and by Conselho Nacional de Desenvolvimento Científico e Tecnológico (CNPq) - Chamada ConhecBrasil-REDES - project number 444683/2024-0. We are in debt with Dr. Jeffrey Bose at the University of Kansas Medical Center for providing strain JE2 and assistance with the animal experiment protocol. We also thank Dr. Chamani Perera, Dr. Srilaxmi Mali Patel and Indeewara Munasinghe from the Synthetic Chemical Biology Core Facility (SCBC) at the University at University of Kansas (KU) for synthesizing valyl-L-proline, methionylserine and lysopine, provide initial assistance with HPLC work and LC-MS data. We thank Dr. Justin Douglas, Dr. Laurie Harned and Dr. Sarah Ann Neuenswander from NMR core facility, University of Kansas for analyzing the NMR sample and providing 1-D and 2-D NMR data.

## Ethical Approval

Animal studies were conducted in strict accordance with approved protocols (293-01) by the University of Kansas Institutional Animal Care and Use Committee.

## Author contributions

A.S. and R.B.R.F. conceptualized the study and wrote the paper. A.S. performed purification and identification experiments, K.H. performed animal and cell culture experiments, W.R.R. performed animal experiments. A.C.S.C.O and L.C.M.A provided critical scientific input to the experimental design. R.B.R.F. acquired the funding and supervised the study. All authors read and approved the final manuscript.

## Supplementary Information

**Figure S1.** Impact of different concentrations of a racemic mixture of pyroglutamic acid (PCA) against *S. aureus* strain JE2 biofilm formation. Mean values of 3 independent experiments are shown, and error bars represent the SD. Statistical analyses were performed with the Kruskal-Wallis test. \*\*\*\**P*<0.00001; ns: not significant.

**Figure S2.** Impact of additional molecules identified in active fractions of *S. epidermidis* cell-free conditioned media (CFCM) against *S. aureus* biofilm formation. A. Biofilm formation of *S. aureus* growing in the presence of different concentrations of L-glutamyl-L-leucine. B. Biofilm formation of *S. aureus* growing in the presence of different concentrations of L-glutamyl-L-tyrosine. C. Biofilm formation of *S. aureus* growing in the presence of different concentrations of methionylserine. D. Biofilm formation of *S. aureus* growing in the presence of different concentrations of lysopine. Mean values of 3-9 replicates are shown, and error bars represent the SD. Statistical analyses were performed with the Kruskal-Wallis test. * *P*<0.05; \*\*\*\**P*<0.00001; ns: not significant.

**Figure S3.** ^13^C Nuclear Magnetic Resonance (NMR) of a pyroglutamic acid (PCA) standard and HILIC fraction 5 at 500 MHz in D_2_O. A. Pyroglutamic acid standard. B. HILIC fraction 5.

**Figure S4.** Correlation Spectroscopy (COSY) ^1^H-^1^H interaction at 500 MHz in D_2_O. A. Pyroglutamic acid standard. B. HILIC fraction 5.

**Figure S5.** Heteronuclear Single Quantum Coherence (HSQC) ^1^H-^13^C interaction at 500 MHz in D_2_O. A. Pyroglutamic acid standard. B. HILIC fraction 5.

**Figure S6.** Heteronuclear Multiple Bond Correlation (HMBC) ^1^H-^13^C interaction at 500 MHz in D_2_O. A. Pyroglutamic acid standard. B. HILIC fraction 5.

**Figure S7.** Liquid chromatography-mass spectrometry (LC-MS) analysis of compounds. A. Pyroglutamic acid standard. B. HILIC Fraction 5.

**Figure S8.** Impact of pyroglutamic acid (PCA) on A549 viability analyzed through MTT ((3- (4,5-dimethylthiazol-2-yl)-2,5-diphenyltetrazolium bromide) assay.

