## Supplementary material for "A microbiome-derived metabolite from *Staphylococcus epidermidis* inhibits *Staphylococcus aureus* biofilm formation and virulence": Suplemental Information

### Supplementary Information

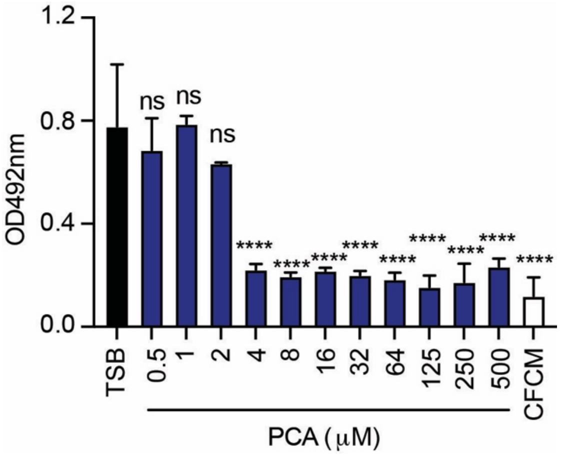


**Figure S1.** Impact of different concentrations of a racemic mixture of pyroglutamic acid (PCA) against *S. aureus* strain JE2 biofilm formation. Mean values of 3 independent experiments are shown, and error bars represent the SD. Statistical analyses were performed with the Kruskal-Wallis test. ****$P$<0.00001; ns: not significant.

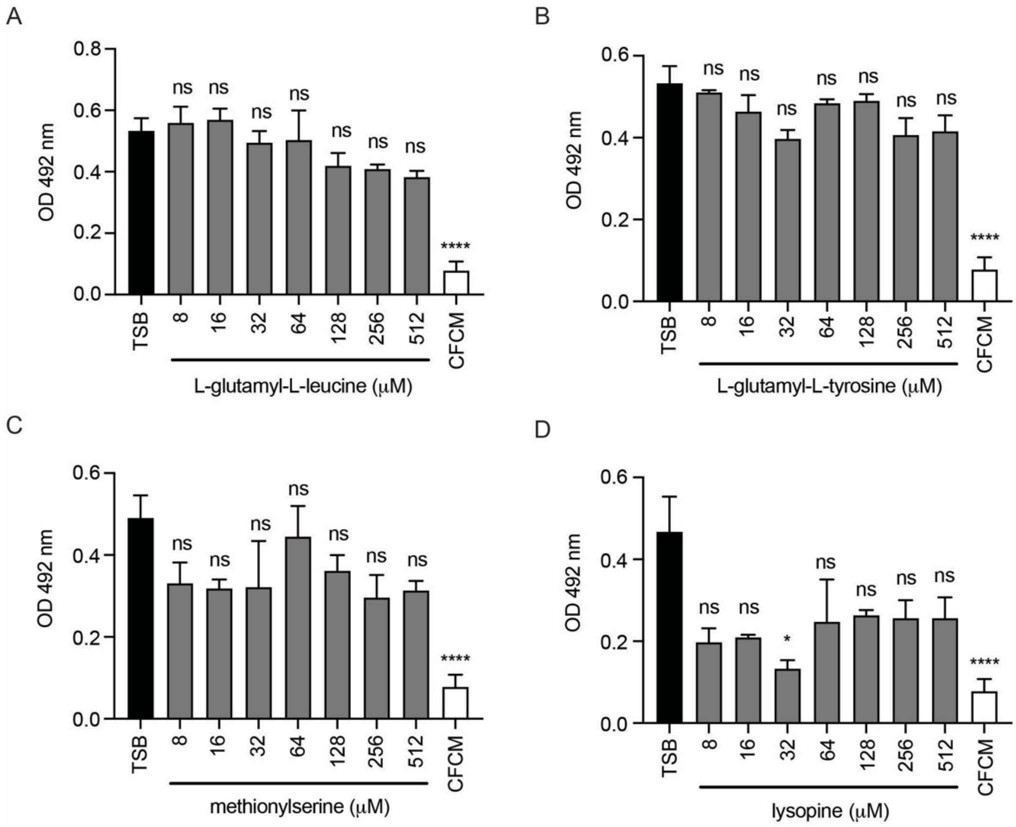


**Figure S2**. Impact of additional molecules identified in active fractions of *S. epidermidis* cell-free conditioned media (CFCM) against *S. aureus* biofilm formation. A. Biofilm formation of *S. aureus* growing in the presence of different concentrations of L-glutamyl-L-leucine. B. Biofilm formation of *S. aureus* growing in the presence of different concentrations of L-glutamyl-L-tyrosine. C. Biofilm formation of *S. aureus* growing in the presence of different concentrations of methionylserine. D. Biofilm formation of *S. aureus* growing in the presence of different concentrations of lysopine. Mean values of 3-9 replicates are shown, and error bars represent the SD. Statistical analyses were performed with the Kruskal-Wallis test. *$P$<0.05; ****$P$<0.00001; ns: not significant.

A

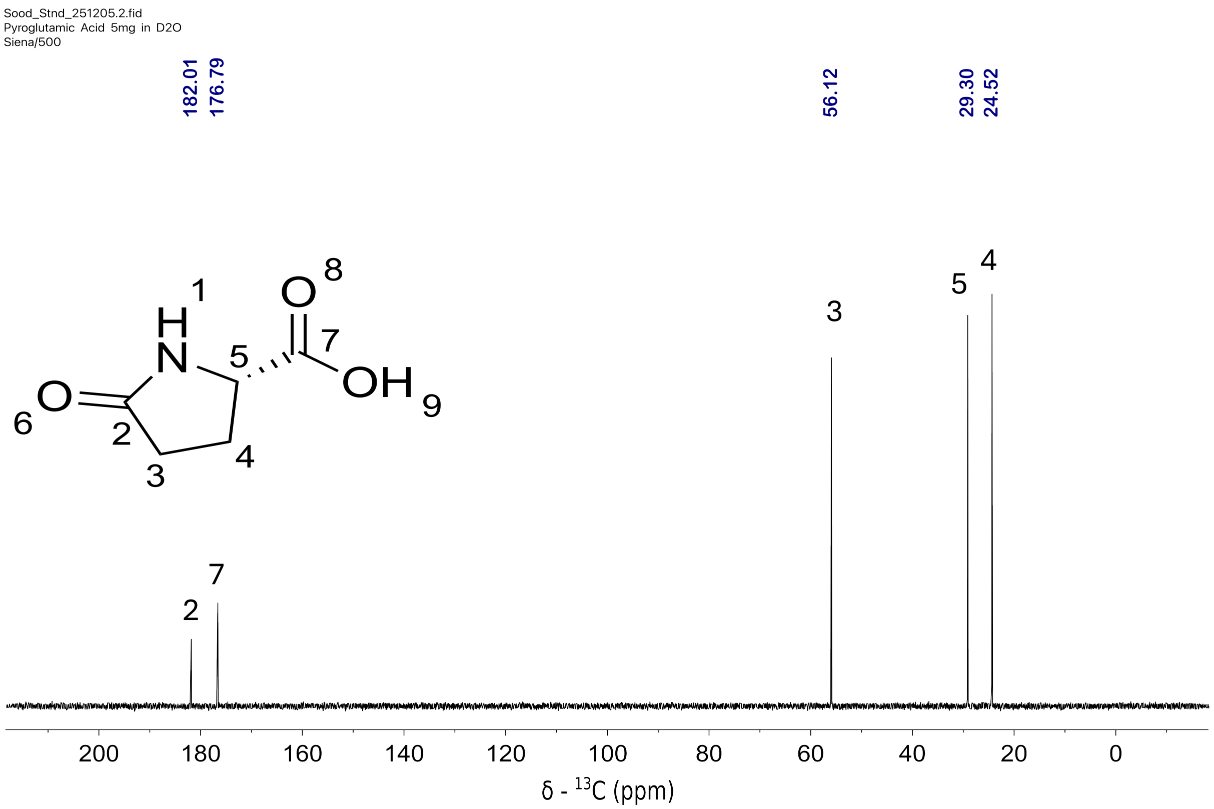


B

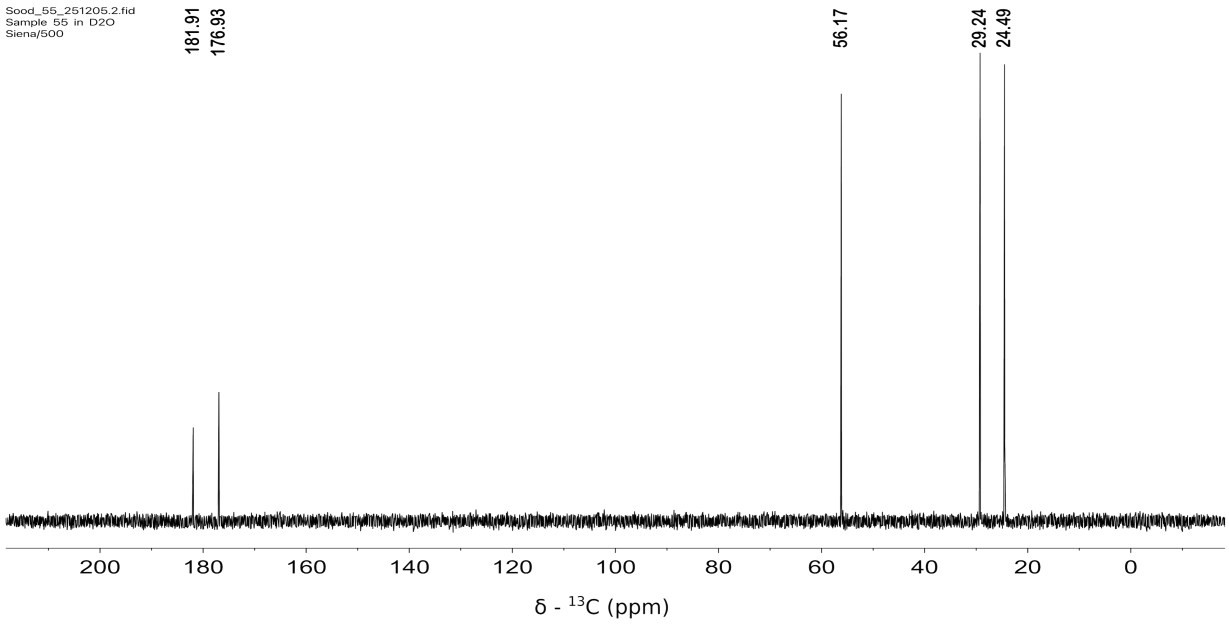


**Figure S3**. $^{13}C$ Nuclear Magnetic Ressonance (NMR) of a pyroglutamic acid (PCA) standard and HILIC fraction 5 at 500 MHz in $D_2O$. A. Pyroglutamic acid standard B. HILIC fraction 5.

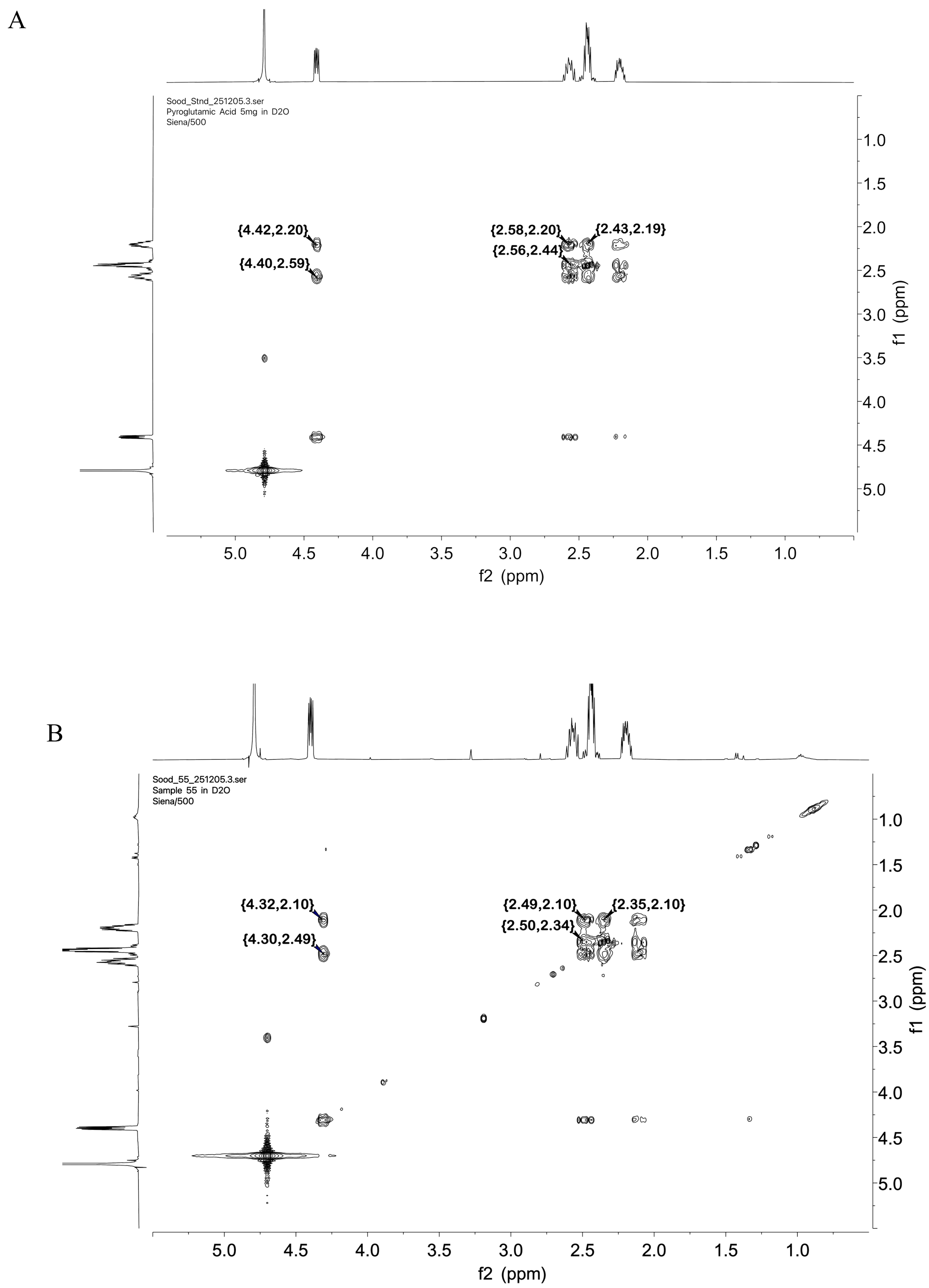


**Figure S4**. Correlation Spectroscopy (COSY) $^1H$-$^1H$ interaction at 500 MHz in $D_2O$. A. Pyroglutamic acid standard. B. HILIC fraction 5.

A

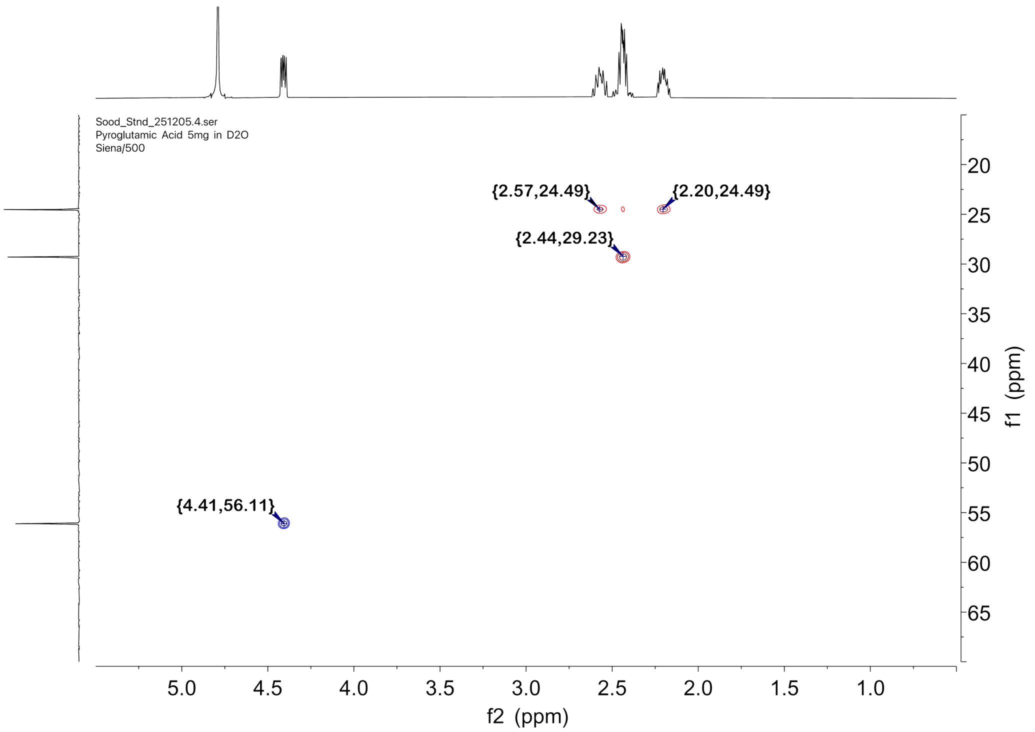


B

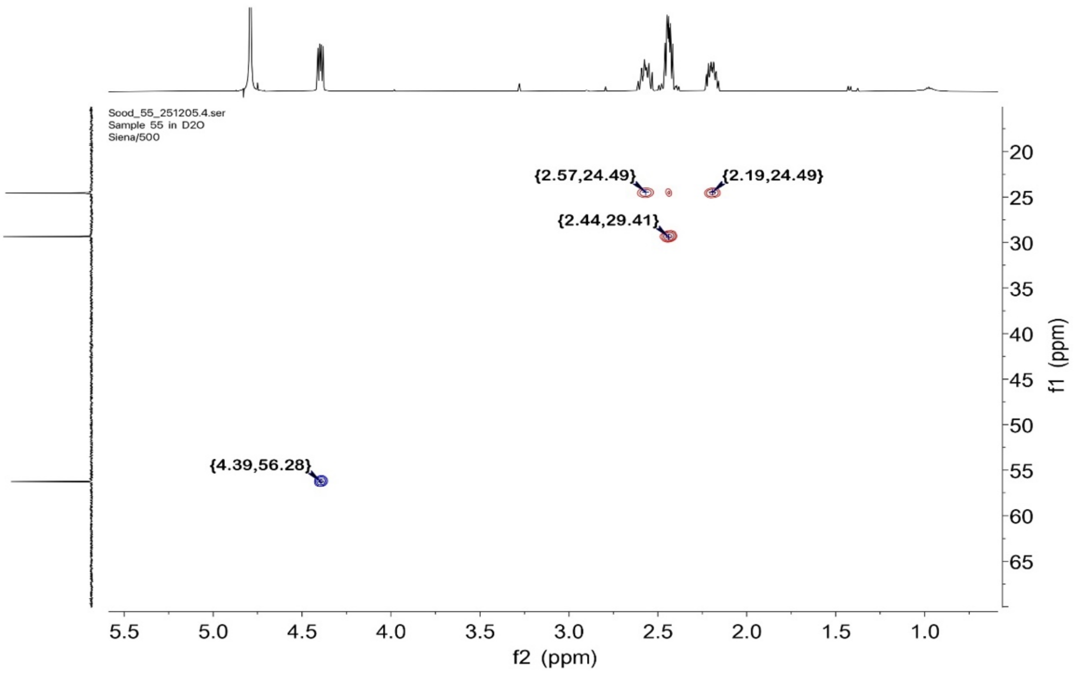


**Figure S5**. Heteronuclear Single Quantum Coherence (HSQC) $^{1}H$-$^{13}C$ interaction at 500 MHz in $D_2O$. A. Pyroglutamic acid standard. B. HILIC fraction 5.

A

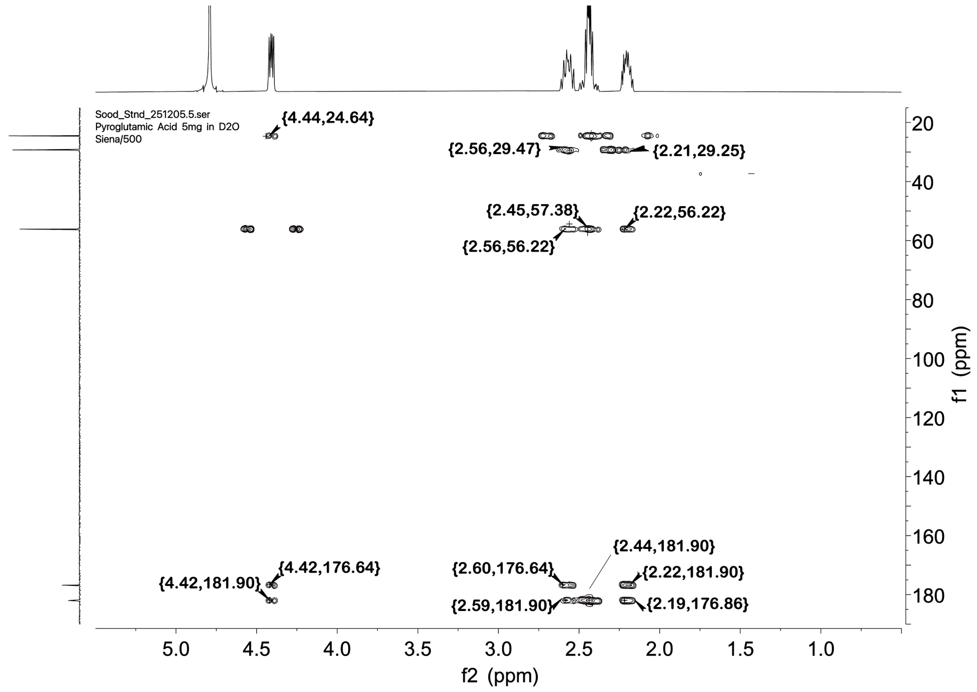


B

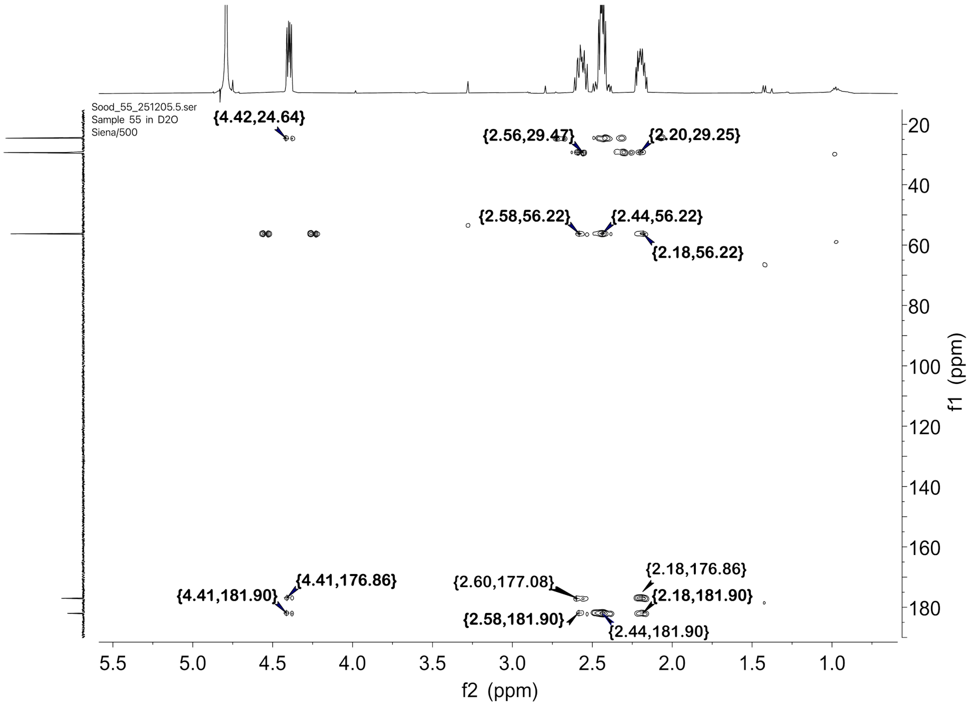


**Figure S6**. Heteronuclear Multiple Bond Correlation (HMBC) $^{1}H$-$^{13}C$ interaction at 500 MHz in $D_2O$. A. Pyroglutamic acid standard. B. HILIC fraction 5.

A

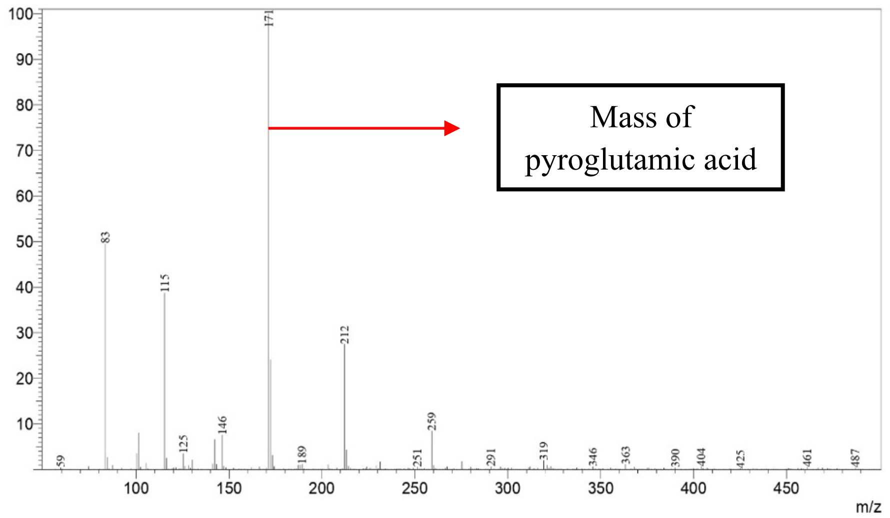


B

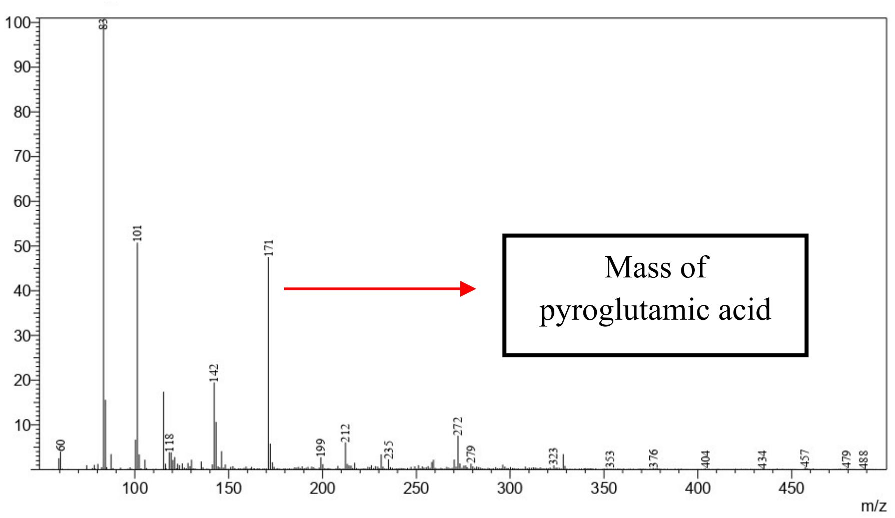


**Figure S7**. Liquid chromatography-mass spectrometry (LC-MS) analysis of compounds. A. Pyroglutamic acid standard. B. HILIC Fraction 5.

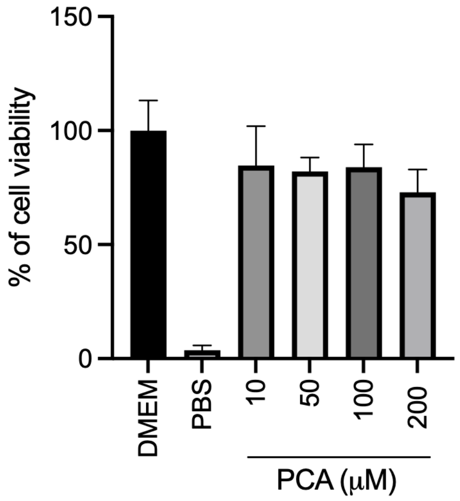


**Figure S8**. Impact of pyroglutamic acid (PCA) on A549 viability analyzed through MTT ((3-(4,5-dimethylthiazol-2-yl)-2,5-diphenyltetrazolium bromide) assay.

Table S1. Metabolites enriched in bioactive fractions

| Name | Formula | *m/z* | RT [min] | Log2 Fold change of active fractions compared to inactive fraction 19 | | | | Log2 Fold change of active fractions compared to inactive fraction 26 | | | |
|---|---|---|---|---|---|---|---|---|---|---|---|
| | | | | 2C | 2D | 2E | 2F | 2C | 2D | 2E | 2F |
| (8E)-2-amino-8-octadecene-1,3,4-triol | C18 H37 N O3 | 338.2658 | 7.976 | 4.47 | 8.1 | 1.15 | 8.22 | 4.8 | 8.43 | 1.48 | 8.56 |
| methionylserine | C8 H16 N2 O4 S | 237.08993 | 1.548 | 10.79 | 11.46 | 2.68 | 5.94 | 2.74 | 3.41 | -5.37 | -2.11 |
| pyroglutamic acid | C5 H7 N O3 | 130.04983 | 3.257 | 2.67 | 4.07 | 1.92 | 4.54 | 3.08 | 4.47 | 2.32 | 4.94 |
| γ-glutamyl-leucine | C11 H20 N2 O5 | 261.14375 | 3.572 | 11.99 | 9.73 | 3.31 | 4.13 | 13.04 | 10.78 | 4.36 | 5.18 |
| lysopine | C9 H18 N2 O4 | 219.13365 | 2.224 | 8.28 | 9.73 | 9.47 | 3.72 | 8.83 | 10.29 | 10.02 | 4.28 |
| γ-glutamyl-leucine | C11 H20 N2 O5 | 261.14381 | 3.441 | 2.04 | 3.27 | 11.31 | 10.42 | 3.25 | 4.48 | 12.52 | 11.63 |
| mebutamate* | C10 H20 N2 O4 | 233.1492 | 3.21 | 11.08 | 12.1 | 3.97 | 1.86 | 11.76 | 12.78 | 4.65 | 2.54 |
| valyl-proline | C10 H18 N2 O3 | 215.13861 | 3.458 | 12.35 | 12.8 | 3.1 | 1.83 | 12.86 | 13.31 | 3.61 | 2.34 |
| N6-acetyl-N6-hydroxy-lysine | C8 H16 N2 O4 | 205.11806 | 1.917 | 1.67 | 4.77 | 8.48 | 1.45 | 2.43 | 5.53 | 9.24 | 2.21 |
| γ-glutamyl-tyrosine | C14 H18 N2 O6 | 311.12309 | 3.161 | 8.96 | 8.8 | 1.1 | 1.42 | 8.98 | 8.82 | 1.12 | 1.44 |
| uracil | C4 H4 N2 O2 | 113.03467 | 2.009 | 8.81 | 2.76 | 1.02 | 1.09 | 9.23 | 3.18 | 1.44 | 1.51 |

RT: retention time; *: not naturally found.; shaded areas: compound enriched over 2.3 log2 fold (5 fold) in comparison with inactive fractions.
